# Orally active mGluR2/3 metabotropic antagonist pro-drug mimics the beneficial effects of physical exercise on neurogenesis, behavior, and exercise-related molecular pathways in an Alzheimer’s disease mouse model

**DOI:** 10.1101/2022.01.31.477906

**Authors:** Georgina Perez-Garcia, Mesude Bicak, Jacqueline Buros, Jean-Vianney Haure-Mirande, Gissel M. Perez, Alena Otero-Pagan, Miguel A. Gama Sosa, Rita De Gasperi, Mary Sano, Fred H. Gage, Carrolee Barlow, Joel T. Dudley, Benjamin S. Glicksberg, Yanzhuang Wang, Benjamin Readhead, Michelle E. Ehrlich, Gregory A. Elder, Sam Gandy

**Author notes:** Perez-Garcia & Bicak are co-first authors. To whom correspondence should be addressed at or.

## Abstract

Modulation of physical activity represents an important intervention that may delay, slow, or prevent mild cognitive impairment (MCI) or dementia due to Alzheimer’s disease (AD). One mechanism proposed to underlie the beneficial effect of physical exercise involves the apparent stimulation of adult hippocampal neurogenesis (AHN). BCI-838 is a pro-drug whose active metabolite BCI-632 is an antagonist at the group II metabotropic glutamate receptor (mGluR2/3). We previously demonstrated that administration of BCI-838 to a mouse model of cerebrovascular accumulation of oligomeric Aβ^E22Q^ (*APP*^*E693Q*^ *= “Dutch APP”*) reduced learning behavior impairment and anxiety, both of which are associated with the phenotype of the *Dutch APP* mice. Here we show that (i) administration of BCI-838, physical exercise, or a combination of BCI-838 and physical exercise enhanced AHN in a four-month old mouse model of AD amyloid pathology (*APP*^*KM670/671NL*^*/ PSEN1*^*Δexon9*^ = APP/PS1), (ii) administration of BCI-838 alone or associated with physical exercise led to stimulation of AHN and improvement in both spatial and recognition memory, (iii) significantly, the hippocampal dentate gyrus transcriptome of APP/PS1 mice following BCI-838 treatment up-regulated brain-derived neurotrophic factor (BDNF), PIK3C2A of the PI3K-MTOR pathway, and metabotropic glutamate receptors, and down-regulated EIF5A of ketamine-modulating mTOR activity, and finally (iv) qPCR findings validate a significantly strong association between increased BDNF levels and BCI-838 treatment. Our study points to BCI-838 as a safe and orally active compound capable of mimicking the beneficial effect of exercise on AHN, learning behavior, and anxiety in a mouse model of AD neuropathology.

## Introduction

Physical exercise (PE) has important neuroprotective and pro-cognitive benefits^1, 2^. One long-suggested mechanism underlying the beneficial effects of PE on central nervous system (CNS) function is its effect on neurogenesis in the adult hippocampal dentate gyrus (DG) and the subventricular zone (SVZ) of the lateral ventricles^3, 4^. In these regions, new neurons are generated and incorporated into existing neuronal circuits where they promote structural and functional plasticity^4^. Stimulation of adult hippocampal neurogenesis (AHN) has been proposed as a central mechanism of adult brain plasticity and a potential therapeutic target in a variety of psychiatric and neurodegenerative conditions^4^. Drugs such as antidepressants that affect monoaminergic systems, consistently increase AHN^5^. The DG, in particular, is critical for learning and memory, and altered AHN has been implicated in the pathogenesis of neurodegenerative diseases such as Alzheimer’s disease (AD)^6, 7^. A recent review of AHN in neurological diseases^8^ highlighted a key study of the DG from postmortem samples of patients who died from neurodegenerative disorders; the study demonstrated that functions of the neurogenic niche shifted and the cells produced were abnormal in shape and differentiation, further emphasizing the neuronal plasticity characteristic of the hippocampus and its importance to potentially reveal therapeutic targets^9^. Furthermore, a recent study demonstrated that greater physical activity relates to higher levels of presynaptic proteins, suggesting maintenance or building brain resilience^10^.

Exercise-induced AHN has been extensively studied in rodents, and voluntary PE (e.g., running wheel) has been shown to enhance AHN in the DG^11^. Studies utilizing cell fate tracers, e.g., bromodeoxyuridine (BrdU), have shown that exercise increases the proliferation of progenitor cells in the subgranular zone and promotes their survival and differentiation into mature neurons^2, 12^. Increased expression of brain-derived neurotrophic factor (BDNF) may be one mechanism underlying the proneurogenic effect of PE^13^.

Altered AHN has been reported in postmortem brain from humans suffering from AD^6, 7^. In humans, long-term exercise improves blood flow while also increasing hippocampal volume and neurogenesis in subjects with AD^14, 15^. AHN is also altered in transgenic mouse models of AD^16-27^, and exercise has been reported to improve learning behavior in some of these models. For example, in *APP*^*KM670/671NL*^*/PSEN1*^*Δexon9*^ (APP/PS1) transgenic mice, two days of treadmill running for 30 min was associated with increased brain activity, as determined by increased theta rhythm on electroencephalography (EEG). In the same APP/PS1 model, 20 weeks of treadmill training improved learning behavior and reduced brain levels of Aβ42^28^. In the 5xFAD mouse model of AD, four months of voluntary PE in a running wheel improved learning behavior and increased levels of BDNF and synaptic markers in the hippocampus^29^.

BCI-838 (MGS0210) is a group II metabotropic glutamate receptor (mGluR2/3) antagonist pro-drug^30, 31^. BCI-838 is metabolized in the liver into BCI-632, which is the active compound delivered to the brain. As a class of drug, mGluR2/3 receptor antagonists are proneurogenic, as evidenced by their stimulation of AHN while also enhancing learning behavior and exerting broad anxiolytic and antidepressant properties^32^. Previously, BCI-838 administration was observed to improve learning behavior and reduce anxiety in the transgenic mouse model of cerebral amyloidosis^28^ and in a rat model of blast-related traumatic brain injury^31^.

Since both BCI-838 and PE are associated with enhanced AHN, we sought to determine whether BCI-838 might substitute for PE, and whether a combined treatment with both drug and PE might be synergistic. The purpose of this study, therefore, was to investigate whether the administration of BCI-838 might mimic PE and whether combination of BCI-838 with PE might show synergistic or additive effects. Our findings demonstrate that BCI-838 stimulates hippocampal neurogenesis, improves learning behavior, and upregulates BDNF and PIK3C2A levels of MTOR pathway, as well as metabotropic glutamate receptors in APP/PS1 transgenic mice, thereby mimicking the effects of PE.

## Results

### Experimental design for exposure of APP/PS1 mice to PE and treatment with BCI-838

Mouse groups received treatment with BCI-838 or vehicle for one month with or without PE (Fig. 1). PE consisted of *ad libitum* access to a running wheel. Three-month-old mice were divided into five experimental groups: (1) wild type (WT) mice treated with vehicle; (2) APP/PS1 mice treated with vehicle (APP/PS1 control); (3) APP/PS1 mice treated with vehicle and exposed to running wheels (APP/PS1 + PE); (4) APP/PS1 mice treated with 5 mg/kg BCI-838 (APP/PS1 + BCI-838); and (5) APP/PS1 mice treated with 5 mg/kg BCI-838 and exposed to running wheels (APP/PS1 combination) (Fig. 1a). Three cohorts of mice were studied. One cohort was used for behavioral analysis, the second for neurogenesis assays and the third for transcriptomic analysis. Spontaneous running wheel activity was analyzed in all cohorts. When compared to APP/PS1 control, APP/PS1 + BCI-838 showed decreased spontaneous activity during the last two weeks of the treatment protocol, as evidenced by reduced counts of running wheel activity (Fig. 1b).

**Fig 1.**
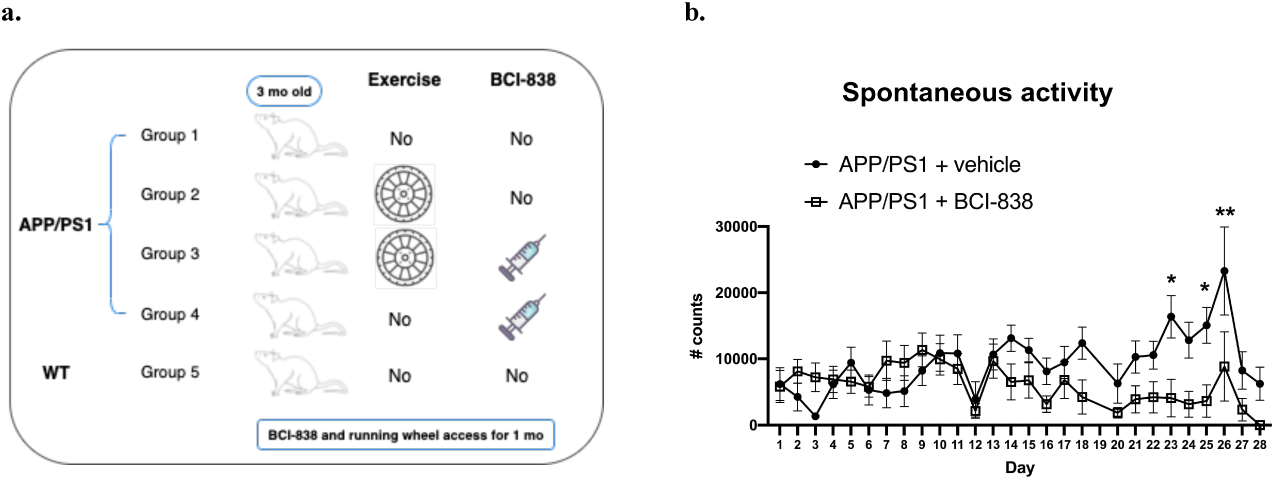
Experimental design and spontaneous activity. **a** APP/PS1 mice were divided into four groups: Group 1 no exercise / no drug; Group 2 exercise / no drug; Group 3 exercise + drug; and Group 4 no exercise /drug. Group 5 consisted of WT mice (C57BL6/J). Groups receiving exercise were given *ad libitum* access to a running wheel and treatment with drug or vehicle was continued for one month. Two cohorts of mice were used. One cohort was used for behavioral analysis whereas a second was sacrificed for neurogenesis and biochemical studies. **b** BCI-838 decreased spontaneous activity after two weeks of treatment. Numbers of running wheel turns (# counts) are shown. APP/PS1 mice treated with BCI-838 decreased running wheel use during the last two weeks of drug treatment and running wheel exposure. Repeated measures ANOVA over the entire 28 days of treatment revealed a significant difference in running wheel activity within groups (F _3.819, 61.111_ = 3.695, p = 0.011, day*condition p = 0.052 using the Greenhouse-Geisser correction). A test of between subject effects over the 28 days revealed no significant effect of condition (F _1, 16_ = 2.399, p = 0.14). Tests of within subject effects revealed significant effects of running wheel activity if analyzed over 1-14 days (F _3.503, 59.551_ = 3.579, p = 0.014 for activity, F _3.503, 59.551_ = 2.062, p = 0.105 for day*condition) and 14-28 days (F _2.943, 47.094_ = 4.414, p = 0.008 for activity, F _2.943, 47.094_ = 4.414, p = 0.366 for day*condition). There was no difference in running wheel activity between groups over days 1-14 (F _1, 17_ = 0.799) whereas activity was reduced in mice that received exercise + drug between days 14-28 (F _1, 16_ = 6.052, p = 0.026). Values significantly different among groups at individual time points are indicated by asterisks (*p <0.05, **p <0.01, unpaired t-tests). Values are expressed as mean ± SEM.

### Either BCI-838 or PE can improve recognition learning behavior in a novel object recognition test and the Barnes maze

We probed the effect of PE, BCI-838, or the two combined on the learning behavior of APP/PS1 mice (Fig. 2). Novel object recognition (NOR) is a standard behavioral test used to evaluate hippocampus- and perirhinal-dependent learning behavior; it is widely used to assess the progression of learning deficits in mouse models of AD. During training, no differences in object recognition preference were observed among groups (Fig. 2a). During the long-term memory session (LTM, Fig. 2b), WT mice spent more time exploring the novel object (NO) and less time exploring the familiar object (FO) (p = 0.028). APP/PS1 mice explored the FO and the NO the same amount of time (p = 0.989), consistent with the conclusion that APP/PS1 mice at four months of age have impaired learning behavior in this task. However, APP/PS1 mice exposed to one month of PE spent more time exploring the NO when tested 24 h after the training (p = 0.0013). APP/PS1 mice treated with BCI-838 for one month also spent more time exploring the NO (p = 0.0001). APP/PS1 mice treated with the combination of BCI-838 and PE for one month showed a preference for the NO compared to the FO (p = 0.0006). These results suggest that one month of either PE or BCI-838 drug administration, as well as the combination of both, can reverse the impairment of learning behavior in APP/PS1 mice.

**Fig. 2.**
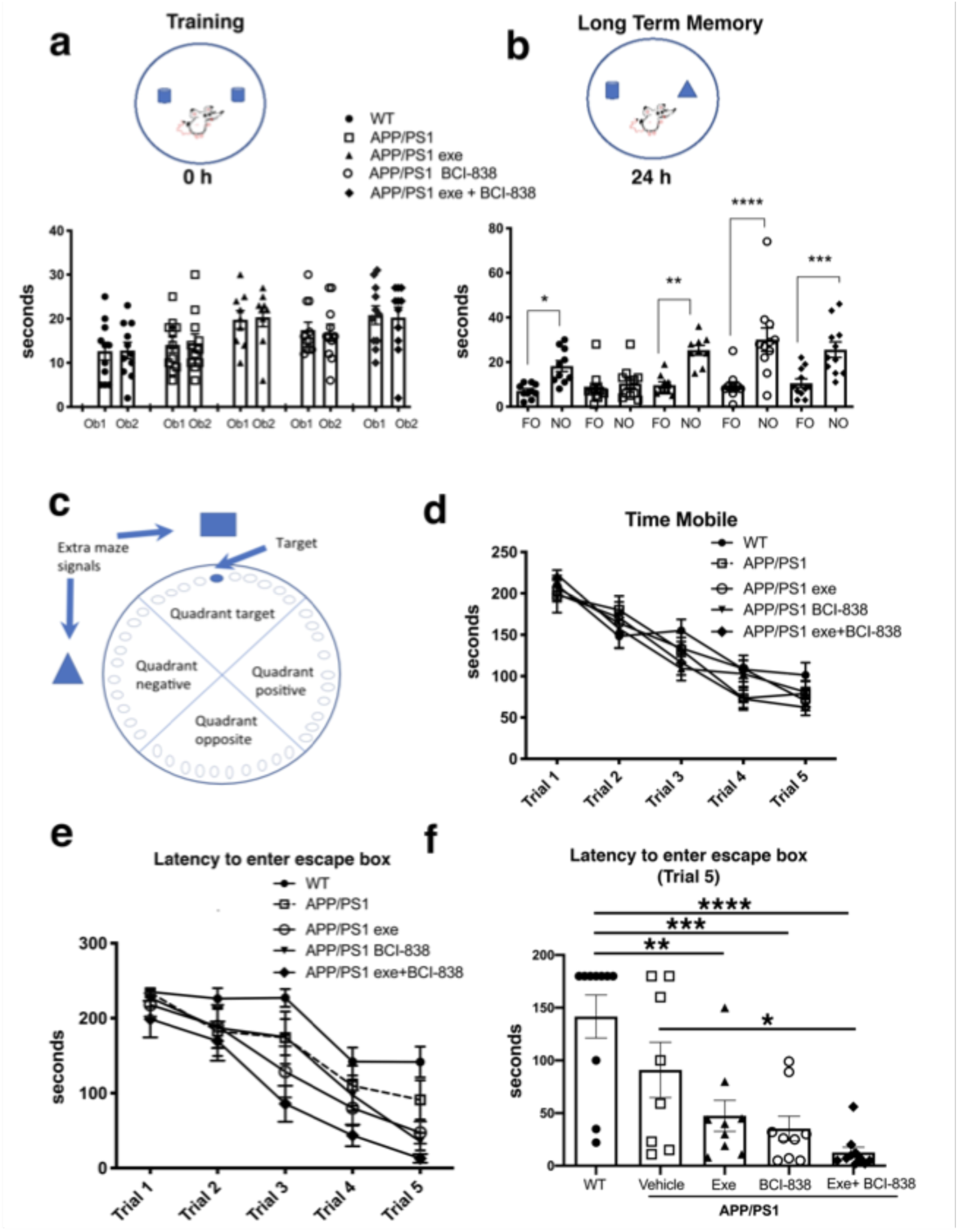
Effect of BCI-838 treatment and exercise on behavioral performance. **a-b** novel object recognition (NOR). **a** Training. No differences were found among groups during the training when the mice explored the similar objects, Object 1 (Ob1) and Object 2 (Ob2). **b** Long-term memory session (LTM). During the LTM testing conducted at 24 h after the training, wild type (WT) mice spent more time exploring the novel object (NO) compared to the familiar object (FO). APP/PS1 mice explored the FO and the NO the same amount of time whereas APP/PS1 mice exposed to exercise, treated with BCI-838 or the combination for one month showed a preference for the NO compared to the FO. Values significantly different are indicated by asterisks (*p <0.05, **p <0.01, ***p< 0.001, ****p< 0.0001, ANOVA, Sidak, multiple comparisons. Values are expressed as mean ± SEM). **c-f** Barnes maze. APP/PS1 mice treated with BCI-838 and exercise showed improved spatial learning as compared to APP/PS1 that did not receive BCI-838 or exercise. **c** Orientation and signals in the Barnes maze. **d** Time in motion over the five trials is shown. Mobility consistently decreased across the trials. A repeated measures ANOVA revealed a significant within-subjects effect (F _4, 184_ = 70.598, p < 0.001) for mobility but no effect of trial*condition (F _16, 184_ = 1.049, p = 0.408). A test of between subject effects revealed no significant group differences (F _4, 46_ = 1.381, p = 0.255). A one-way ANOVA of the day 1 trial also revealed no differences between groups (F _4, 47_ = 0.608, p = 0.659). **e** Latency to enter the escape box. Tests of within subject effects showed that all groups demonstrated a progressive decrease in latency to enter the escape box (repeated measures ANOVA, F _3.385, 148.961_ = 10.688, p < 0.001, for latency, F _3.385, 148.961_ = 13.542, p = 0.671 for trial*condition, Greenhouse-Geisser correction). There were, however, significant between-subject effects (F _4, 44_ = 4.198, p = 0.006). **f** A one-way ANOVA of trial 5, revealed significant between-group effects (F _4, 41_ = 10.09, p < 0.0001). Values significantly different among groups are indicated by asterisks, *p <0.05, **p <0.01, *** p < 0.001, **** p < 0.0001 (Sidak multiple comparisons). Values are expressed as mean ± SEM.

We next tested mice in a Barnes maze, a behavioral test used to probe hippocampus-dependent spatial learning behavior. In this task, mice use visual cues to learn the location of an escape box (Fig. 2c). Mobility was not altered between groups (Fig. 2d). In the training phase, the latencies of all groups decreased across the five trials, indicating learning of the task (Fig. 2e). However, by the fifth trial, APP/PS1 mice treated with PE, with BCI-838, or with the combination exhibited decreased latencies to enter the escape box, as compared to WT mice treated with vehicle (Fig. 2f). Mice treated with the combination exhibited shorter latencies than APP/PS1 mice treated with vehicle (Fig. 2f). APP/PS1 mice treated with vehicle did not differ from vehicle-treated WT mice. Taken together, these results show that the administration of BCI-838, PE or the combination can reverse cognitive deficits or enhance cognitive function in APP/PS1 mice.

### BCI-838 administration or PE, alone or in combination, enhances hippocampal neurogenesis

We next probed the effect of PE, BCI-838 administration, or a combination of both on AHN in APP/PS1 mice (Fig. 3). Using immunofluorescence staining, we quantified newly generated neurons (doublecortin-labeled; DCX) cells; Fig. 3a-e) and newly generated cells (BrdU-labeled; Fig. 3f) in APP/PS1 mice treated with vehicle or PE, BCI-838, or both. While the number of DCX-labeled neurons was unchanged in APP/PS1 mice treated with vehicle vs. WT mice, this number was increased in the APP/PS1 mice treated with BCI-838 or with BCI-838 plus PE, when either was compared to APP/PS1 treated with vehicle (F _4, 16_ = 1.367, p = 0.0025; p < 0.05, Sidak post hoc test, Fig. 3a-e, g). Moreover, we observed that the total number of BrdU-labeled cells was increased in APP/PS1 mice treated with BCI-838 or BCI-838 plus PE as compared to APP/PS1 mice treated with vehicle (F _4, 14_ = 1.734; p = 0.0148, p < 0.05, Fig. 3f, h).

**Fig. 3.**
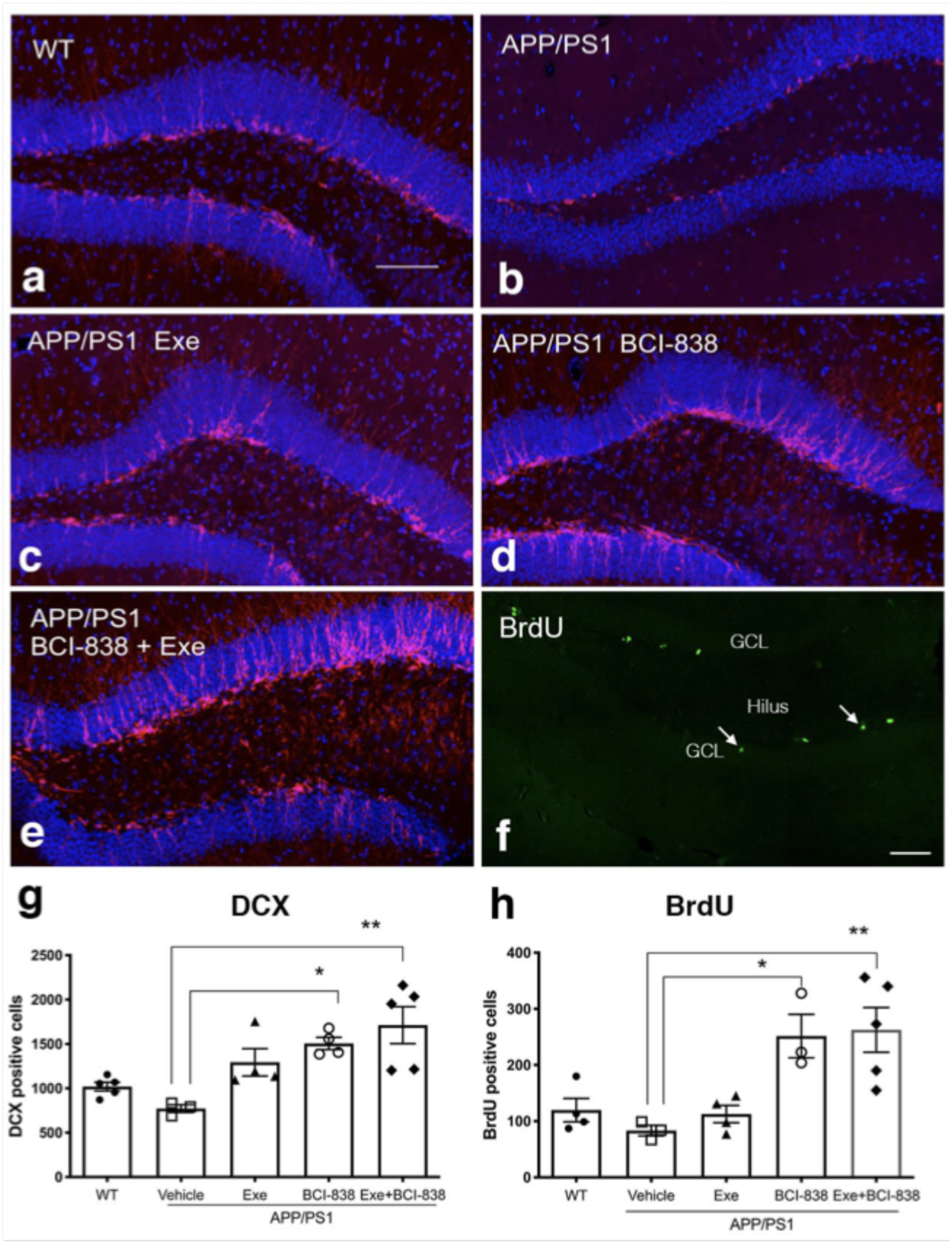
Quantification of neurogenesis in APP/PS1 control, PE, BCI-838 or combination. **a-e** Images of DCX staining in WT, APP/PS1, APP/PS1 + PE, APP/PS1 + BCI-838, and APP/PS1 combination. **f** Representative image of BrdU staining in a WT mouse. The hilus and granule cell layer (GCL) are indicated. Arrows mark a BrdU-labeled cell in the subgranular zone. Scale bar 100 µm. **g** Total number of doublecortin-labeled cells was increased in the APP/PS1 + BCI-838 and APP/PS1 combination as compared to APP/PS1 control. Values are expressed as mean ± SEM and differences among groups are indicated by asterisks (*p < 0.05, **p < 0.01, Sidak multiple comparisons). **h** Total number of BrdU-labeled cells was increased in APP/PS1 + BCI-838 and APP/PS1 combination as compared to APP/PS1 control. Values significantly different among groups are indicated by asterisks as in panel **g**.

### Transcriptomic profiles from the hippocampal DG of APP/PS1 mice show that BCI-838 affects both exercise-related and exercise-independent molecular pathways

We generated transcriptomic profiles from all four groups of four-month-old APP/PS1 and one group of WT mice and performed RNA sequencing on 24 samples from their DG (Fig. 4). When we performed variance partitioning analysis as part of quality control, we noticed a high fraction of unexplained variance in our RNA-Seq dataset and used surrogate variable analysis to address this issue (see Methods and Supplemental Data 1). We then performed differential gene expression analysis to compare all mice models with each other, from APP/PS1 mice exposed to either BCI-838, PE or a combination, as well as WT mice. Differentially expressed genes (DEGs) were identified at a false discovery rate (FDR) of 0.05 (see Supplemental Data 2 for a list of all DEGs for all comparisons). To identify biological pathways that may be differentially dysregulated by administration of BCI-838 or exposure to exercise, we performed gene set enrichment analysis (GSEA) on the identified DEGs, where we identified enriched biological pathways, gene ontology sets, and transcription factor binding sites at an adjusted p-value < 0.05 (see Supplemental Data 3 for GSEA results). Top DEGs and GSEA results sorted by their adjusted p-values are summarized in Figure 5.

**Fig. 4.**
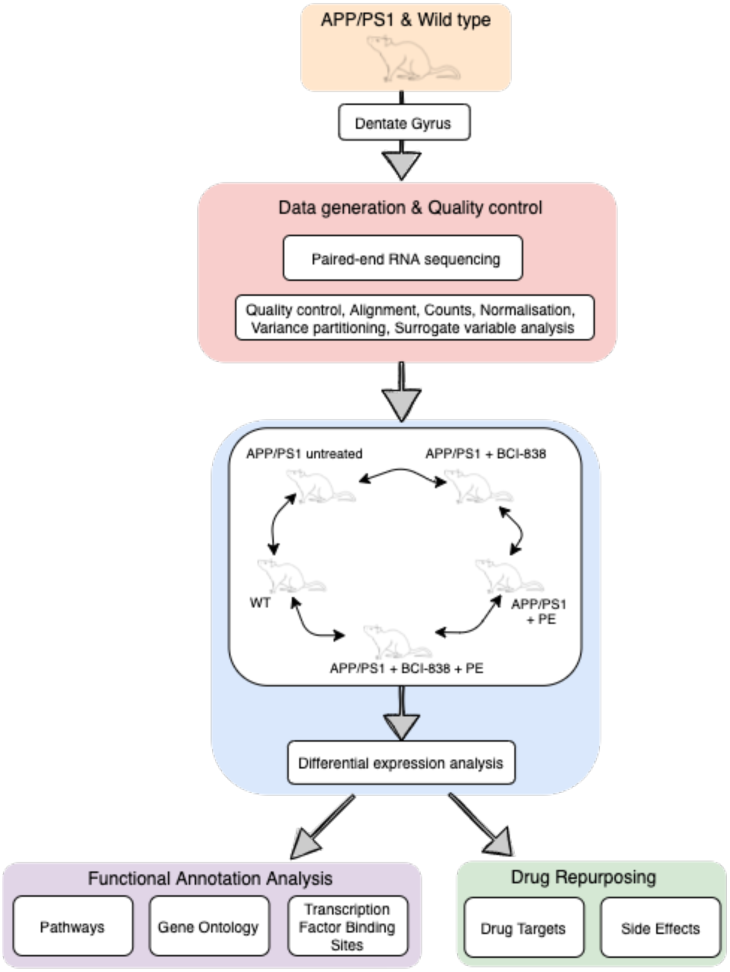
Schematic overview of mouse AD transcriptome analysis. RNA sequencing was performed on dentate gyrus for five groups of four-month-old APP/PS1 and WT mice, comprising a total of 25 samples.

**Fig 5.**
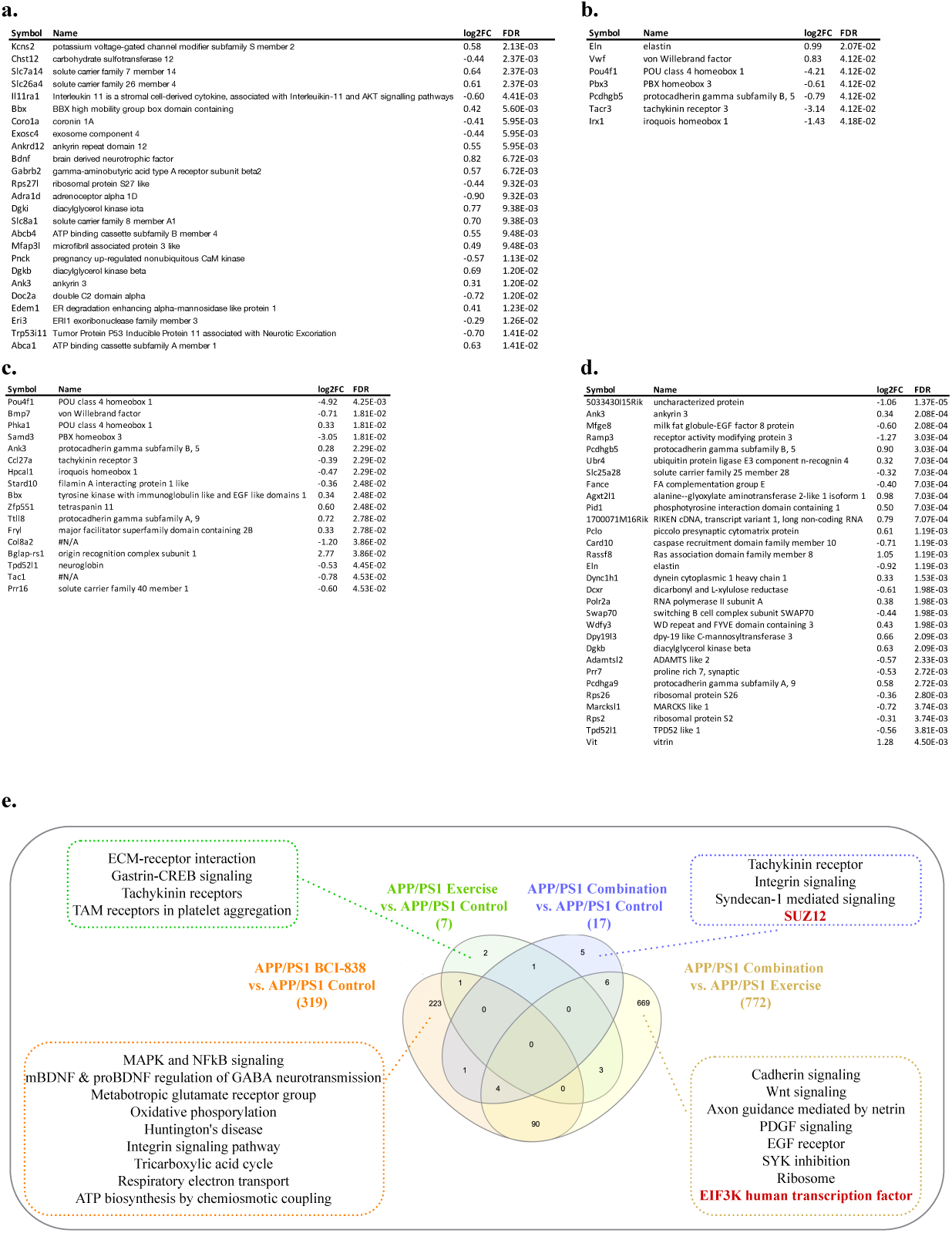
Differential gene expression (DEG) and enrichment analysis summary in dentate gyrus (DG) of APP/PS1 mice treated with BCI-838, exercise or a combination of both. **a** Top DEGs in DG of APP/PS1 mice treated with BCI-838 vs. APP/PS1 treated with vehicle. **b** Top DEGs in DG of APP/PS1 mice treated with exercise vs. APP/PS1 treated with vehicle. **c** Top DEGs in DG of APP/PS1 mice treated with a combination of both BCI-838 and exercise vs. APP/PS1 treated with vehicle. **d** Top DEGs in DG of APP/PS1 mice treated with a combination of both BCI-838 and exercise vs. APP/PS1 treated with exercise. **e** Venn Diagram of DEGs and selected pathway enrichments for known or suspected relevance to AD pathophysiology, shared across different comparisons of APP/PS1 mice groups (transcription factor enrichments shown in red).

The most striking evaluation of all groups are the transcriptomic changes induced by one month of BCI-838 administration in the DG of APP/PS1 (APP/PS1 + BCI-838), which identified 319 DEGs. 191 up-regulated DEGs from this analysis include BDNF, as well as PIK3C2A of the PI3K-MTOR pathway, whereas 128 down-regulated DEGs include IL11RA1, the interleukin 11 receptor of the AKT signaling pathway, as well as EIF5A, the eukaryotic translation initiation factor. This finding is especially notable as it has been shown that the mammalian target of rapamycin (mTOR) pathway is involved in many aspects of neurogenesis by being a chief regulator of cellular energy, nutrient sensor and growth factor inducer via insulin, IGF-1 and BDNF^32, 33^. mTORC1 protein complex controls protein synthesis by phosphorylating downstream targets essential for mRNA translation, 4E-BP1 (eIF-4E binding protein) and ribosome biogenesis^34^. Neuronal growth factors known to support learning and memory such as BDNF and EGF do so through mTOR activation. Furthermore, in relation to the AD brain being an insulin-resistant organ, mTORC1 has been shown to be involved with down-regulating insulin/Akt signaling through an inactivating phosphorylation of IRS-1^35^. Up-regulated DEGs also include ITGB8 of the integrin family, an extracellular matrix (ECM) receptor, known to regulate neurogenesis and neurovascular homeostasis in the adult brain^36^. Of further relevance, dysregulation of several GABA (inhibitory) receptors, GABRB1, GABRB2, and GABRD, and up-regulation of 2 metabotropic glutamate receptor subunits, GRM1 and GRM5, and the ATP-binding AD risk gene ABCA1^37^ are identified among DEGs. GSEA pointed to SP1 as the top transcription factor binding site for down-regulated DEGs; SP1 is known for its participation in the regulation of the mTORC1/P70S6K/S6 signaling^38^. Other differential pathways include neurodegenerative diseases, such as AD and Huntington disease, whereas up-regulated DEGs are found to be enriched for glutamate receptor activity and AKT1 knockdown pathways as expected.

After one month of PE, the transcriptomes of the hippocampal DG of APP/PS1 mice (APP/PS1 + PE) were compared to those of APP/PS1 control mice, leading to the identification of seven DEGs (2 up-regulated, 5 down-regulated) that were found to be enriched for transcripts associated with the response to mTOR inhibitor, with extracellular matrix receptor reaction, with trigeminal nerve development, with peripheral nervous system neuron development, and with polycomb repressive complex component SUZ12, which plays a critical role in regulating neurogenic potential and differentiation of embryonic stem cells^39^. Meanwhile, compared with APP/PS1 control mice, APP/PS1 mice treated with a combination of PE and BCI-838 (APP/PS1 combination) for one month were identified with 17 DEGs (7 up-regulated, 10 down-regulated), also enriched for SUZ12, as well as glycogen metabolism and exercise-induced myalgia pathways.

When APP/PS1 combination was compared with APP/PS1 + BCI-838, we did not observe any DEGs. However, when APP/PS1 combination was compared with APP/PS1 + PE, we identified 772 DEGs, which were found to be enriched in EIF3K and REST transcription factors, both shown to regulate mTOR signaling pathway in adipocyte yeast^38^ and oral cancer cells^41^, respectively. Cadherin signaling, axon guidance mediated by netrin, EGF receptor signaling, SYK inhibition and knockdown were also among differential pathways of relevance with strong associations to mTOR activity and signaling, suggesting neurogenesis and PE, thereby demonstrating the validity of our RNA-Seq data and experiment.

Finally, a comparison of the transcriptomes of the hippocampal DG of APP/PS1 control mice against those of WT mice showed 60 DEGs (41 upregulated, 19 down-regulated) found to be enriched in calcium cation antiporter activity and ATP-dependent microtubule motor activity pathways, in line with highlighted findings from Readhead *et al*.!s transcriptomic analysis of APP/PS1 versus WT^42^.

### BDNF levels in the hippocampal DG of APP/PS1 mice is significantly increased following BCI-838 treatment but not exercise

Increased levels of BDNF have been implicated in the beneficial effects of PE in 5xFAD transgenic mice^29^. Consistent with our DEG findings, where BDNF was only upregulated following BCI-838 treatment and not exercise, qPCR analysis further showed that the BDNF mRNA levels are significantly increased in APP/PS1 mice treated with BCI-838, and not in APP/PS1 mice treated with exercise (Fig. 6a, p = 0.0306, ANOVA, Fisher’s LSD). Furthermore, even though not statistically significant, we observed increased BDNF protein levels in the DG of APP/PS1 mice treated with BCI-838 compared to APP/PS1 mice treated with vehicle or exercise (Fig. 6b). Interestingly, analysis of activity levels showed significantly decreased activity in APP/PS1 mice treated with BCI-838 compared to APP/PS1 mice treated with vehicle (Fig. 6c, p = 0.0353, unpaired t-test). In line with this, despite not significant, correlation analysis demonstrated a higher correlation between increased BDNF mRNA levels and reduced activity levels in APP/PS1 mice treated with BCI-838 (r = 0.447, p = 0.45) compared to APP/PS1 mice treated with vehicle (r = 0.706, p = 0.18) (Fig. 6d).

**Fig. 6.**
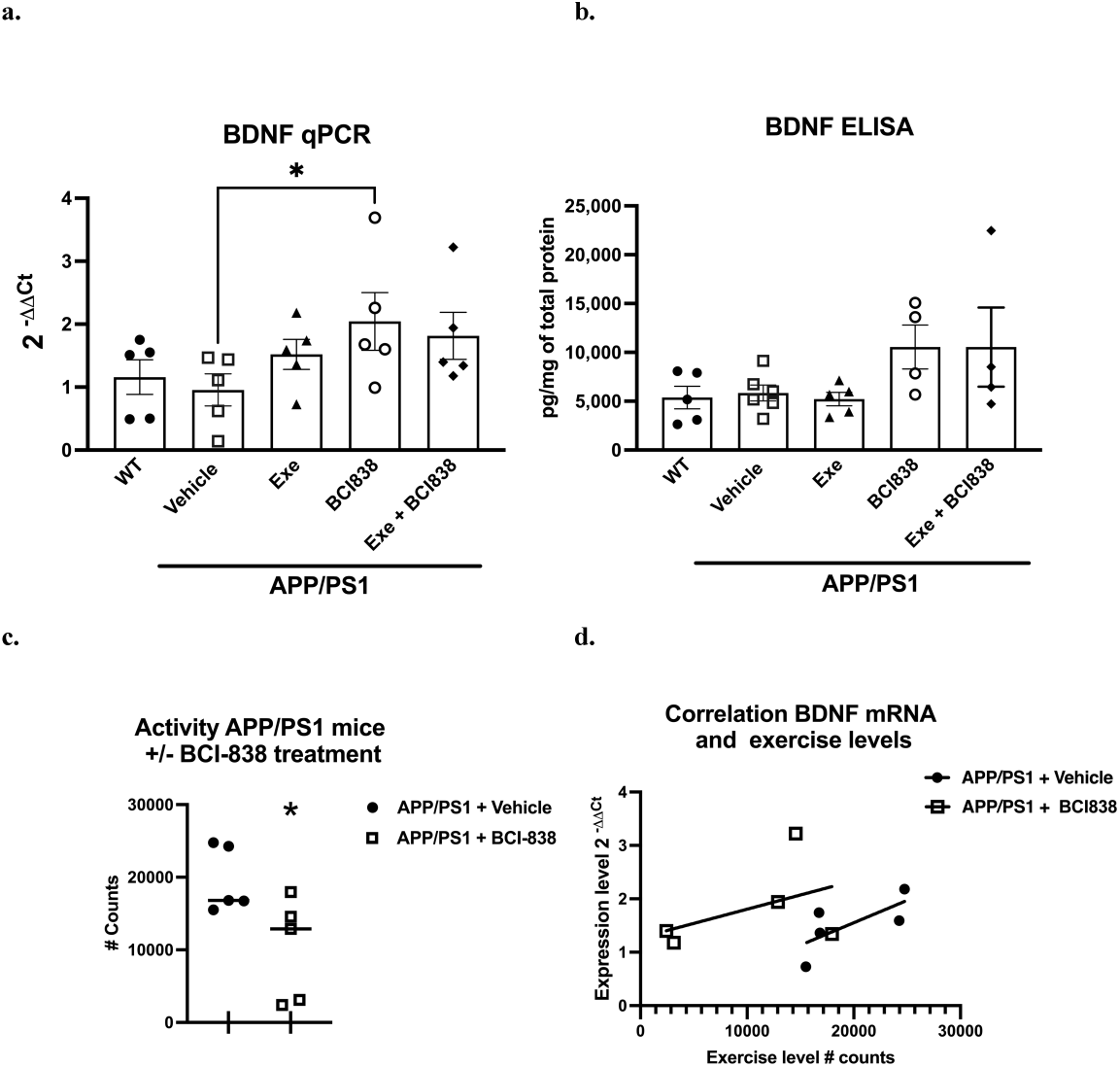
BDNF analysis in APP/PS1 mice treated with BCI-838 or exercise and their correlation with spontaneous activity levels. **a** qPCR for BDNF mRNA in all five groups shows significantly positive correlation with APP/PS1 mice treated with BCI-838 (p = 0.0306, ANOVA, Fisher’s LSD). **b** BDNF protein levels quantified by ELISA, despite not significant, shows increased BDNF protein levels following BCI-838 treatment. **c** Biochemical analysis based on spontaneous running wheel shows decreased average number of total counts in APP/PS1 mice treated with BCI-838 compared to APP/PS1 mice treated with vehicle (p = 0.0353, unpaired t-test). **d** Correlation analysis between BDNF mRNA levels and running wheel counts, despite not significant, demonstrates a higher correlation between increased BDNF mRNA levels and reduced activity levels following BCI-838 treatment (r = 0.706, p = 0.18) compared to vehicle (r = 0.447, p = 0.45).

### Transcriptomically similar drug targets to BCI-838 are enriched for glutamate receptor targets, restore adult neurogenesis, and improve hippocampal memory

We performed computational drug repurposing to identify compounds that induce a transcriptomically similar profile to BCI-838 using a modified Connectivity Mapping^43^ approach (Fig. 7a). We then performed a chemogenomic enrichment analysis^44^ on these compounds to identify drug targets and side effects that were enriched among compounds that were transcriptomically similar to BCI-838 (see Supplemental Data 4 for full drug repurposing results). We observed an enrichment for protein targets that included several glutamate receptor targets, such as GRL1A, NMDE3, GRM2, GRM3, and NMD3B. We also found transcriptomic similarity with other drug targets, such as KDM1A, whose inhibition has been shown to restore adult neurogenesis and improve hippocampal memory^45^, and NLRP3, which plays a role in the regulation of inflammation, immune response, and apoptosis^46^. We also observed that drugs that were transcriptomically similar to BCI-838 were enriched for the side effect of “increasing blood ketones” as well as associated with “pain relief” (Fig. 7b). A ketogenic diet has been shown to inhibit the mammalian target of the rapamycin (mTOR) pathway^47^. Glutamatergic neurotransmission plays a major role in pain sensation and transmission from peripheral tissues into the CNS^48^. These observations comport with findings from DEG and pathway analysis and may suggest follow-up experiments to look for ketogenesis among BCI-838-treated mice.

**Fig. 7.**
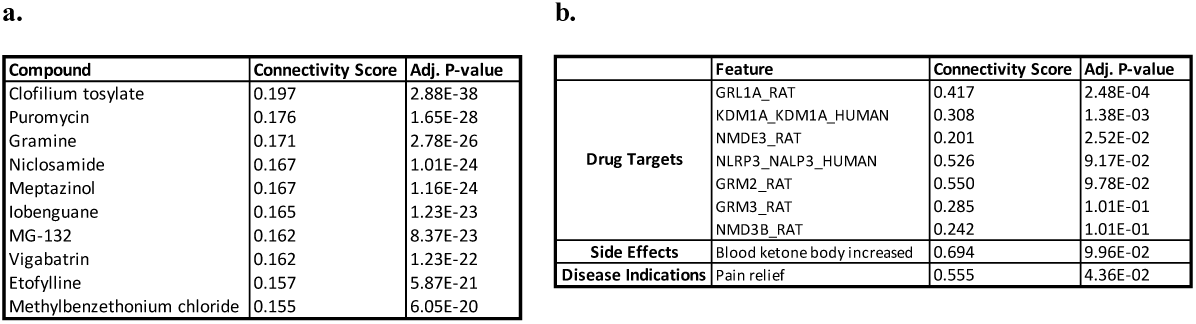
Drug repurposing analysis summary. **a** Top drugs based on transcriptomic similarity. **b** Drug targets and side effects enriched among compounds with transcriptomic similarity to BCI-838.

## Discussion

Metabotropic glutamate receptors (mGluRs) play important roles in regulating glutamatergic neurotransmission. Agents that modulate mGluR2/3 activity have recently gained attention as possible therapeutic agents for a range of mental health-related disorders^49^. BCI-838 is an mGluR2/3 antagonist pro-drug that is metabolized in the liver into the active metabolite BCI-632. We previously reported that BCI-838 administration reduced anxiety and improved learning behavior in the mouse model of accumulation of oligomeric Aβ^E22Q 30^. More recently, we showed that BCI-838 treatment reversed anxiety- and post-traumatic stress disorder-related traits in a rat model of mild blast injury^31^. In both studies, BCI-838 treatment increased brain levels of markers of neurogenesis (BrdU, DCX, PCNA), reduced anxiety-related behaviors, and reversed learning behavior deficits.

PE has important neuroprotective and pro-cognitive effects and has been reported to improve learning behavior in 3XTg-AD mice^50^. Another study using the same APP/PS1 transgenic mouse line studied here revealed that 20 weeks of treadmill training significantly improved learning behavior and reduced brain levels of Aβ42^28^. Like BCI-838, PE stimulated neurogenesis in the hippocampal DG^13^.

Another purpose of the current study was to investigate whether BCI-838 treatment combined with PE could have synergistic effects on learning behavior and AHN in APP/PS1 transgenic mice. Both BCI-838 and PE were associated with restored recognition learning behavior in APP/PS1 mice in a NOR test and improved spatial learning behavior in a Barnes maze. It is interesting how the responses of APP/PS1 mice differed across the two tasks. In NOR, APP/PS1 mice were impaired in recognition memory compared to WT mice and either exercise or drug restored this deficit. By contrast, in the Barnes maze, learning curves did not differ between APP/PS1 mice and WT mice. Thus, the effect of the combined BCI-838 and exercise treatment in the Barnes maze at this early age was more analogous to an enhancement of baseline function rather than to the correction of a learning deficit. The tasks differ in that NOR is dependent on hippocampal and perirhinal function, whereas the Barnes maze is a test that probes hippocampus-dependent spatial learning and memory behavior. Whether differences in the anatomic substrates that support these tasks can explain the differing responses to BCI-838 and exercise remains to be determined.

The differences in performance of APP/PS1 mice in the two tasks may also reflect the relatively young age of the APP/PS1 mice used in this study. A large proportion of studies in transgenic mouse models of AD, including most prior studies in the APP/PS1 mice, have utilized 8-to 12-month-old mice whereas the mice used in this study began treatment at three months of age and were sacrificed at just over four months of age, prior to the significant amyloid deposition in the form of plaques^51^. The failure of the Barnes maze to reveal learning deficits suggests that the cognitive impairment phenotype was probably incompletely developed and that the present study was performed on mice in a state more akin to the human state of mild cognitive impairment (MCI) than to that of fully developed Alzheimer-type dementia. However, given that the AD pathological process is now known to begin decades before symptomatic disease appears^52^, studies of early and late stages of MCI are highly relevant to the human condition, and the current study suggests that BCI-838 and exercise might be most beneficial in the presymptomatic and/or MCI phases of AD.

Alone or in combination, BCI-838 administration and exercise enhanced hippocampal neurogenesis. This conclusion was supported both by BrdU injections given 24 h before sacrifice (which provided a measure of neural progenitor proliferation) and by counts of the numbers of DCX-labeled cells, which provided a measure of the steady-state population of young neurons. Using either measure, BrdU- and DCX-labeled cells were increased following drug treatment or the combination of exercise and drug treatment when compared to APP/PS1 mice treated with vehicle.

Of note, in our studies, neurogenesis was not impaired in APP/PS1 mice treated with vehicle when compared to WT mice. Prior studies in AD transgenic mice have produced inconsistent results concerning the effect of familial AD mutations on AHN^16-27^, including in studies using the APP/PS1 mice studied here^16, 22-26^. In this study, the absence of impaired AHN in APP/PS1 mice compared to WT mice suggests that the effect of BCI-838 and exercise on the neurogenesis may provide enhancement of baseline function rather than restoration of disease-related deficiency in function. This point may be important in determining whether PE intervention is most effective as prophylaxis versus arrest or retardation of symptom progression. Additional studies will be required to determine the fate of newly generated cells and whether BCI-838 and/or PE stimulate the production of new neurons that become integrated into functional neuronal networks as well as whether neurogenesis enhancement is required for the effects of BCI-838 with or without PE.

Why a mGluR2/3 receptor antagonist should mimic the effects of PE is unclear. PE exerts a variety of beneficial effects on neuroplasticity, spatial learning, and memory^52^. Neurogenesis in adult brain is likely regulated through a combination of factors^3^, and PE also acts on multiple pathways^2^. Mechanistically, many studies have emphasized PE-related effects on trophic factor production, especially BDNF, insulin-like growth factor-1 (IGF-1), and vascular endothelial growth factor (VEGF)^53^. A recent study in AD transgenic 5xFAD mice further demonstrated that exercise-induced AHN improved cognition along with increased levels of BDNF^54^.

More recent work has emphasized how exercise affects secreted factors in the periphery^55^. In particular, myokines secreted by muscle during exercise have been identified to improve learning and memory by regulating hippocampal function^56, 57^. Cathepsin B, a PE-related myokine, crosses the blood-brain barrier and enhances BDNF production^57^. Furthermore, irisin, the cleaved and circulating form of the exercise-induced fibronectin type III domain-containing membrane protein (FNDC5), stimulates BDNF production^58, 59^; its genetic deletion is shown to impair cognitive function in exercise, aging and AD; and its peripheral application is found to be sufficient to rescue the cognitive decline in mouse models of AD^58^. In our study, we did not detect differential gene expression of FNDC5 in the DG of the APP/PSEN1 either following BCI-838 administration or after exercise vs. sedentary APP/PSEN1 mice, which may be due to the relatively young age of the APP/PSEN1 mice used (four months) and the absence of neuronal and synaptic loss in this model at this age. Furthermore, the length of voluntary aerobic exercise was also limited (one month) as a consequence of the young aged mice.

Recently, integrins have been identified as receptors for irisin in bone and fat cells^60^. Single-cell RNA-seq data from the murine DG found ITGAV expressed in neuronal and non-neuronal cells and ITGB5 expressed in astrocytes and microglia, raising the possibility that irisin might mediate glia activation through integrin receptor complexes^58^. When we scrutinized our data for possible evidence of transcriptomic changes among integrin family members, we observed upregulation of ITGB8, an ECM receptor that regulates neurogenesis and neurovascular homeostasis in the adult brain^36,^ following BCI-838 administration to *APP/PSEN1* mice, at an FDR of 5%, as well as ITGB6 and ITGB7 at an FDR of 1%.

By contrast, mGluR2/3 receptors function primarily as presynaptic autoreceptors that, when stimulated, inhibit glutamate release^49, 62^. The antidepressant effects of agents acting on the mGlu2/3 receptor have been widely studied in animal models. mGlu2/3 receptor blockade appears to act at least in part through mTOR signaling, which may contribute to the sustained antidepressant-like effects of mGlu2/3 receptor antagonists^58^. However, the full synaptic and neural mechanisms of their action (in particular, how mGluR2/3 antagonists stimulate AHN) remain to be clarified^50, 62^.

In this context, it is interesting to compare our results with recent work in transgenic 5xFAD mice^29^, which revealed that chronic voluntary PE in running wheels for four months, beginning at two months of age, improved learning behavior in 5xFAD mice; meanwhile, our current study reveals the improvement in one month of PE. PE-mediated enhancement of neurogenesis in the hippocampus has been suggested to be mediated through BDNF signaling^64^. In our studies, while a short treatment with BCI-838 or PE was associated with increased neurogenesis in the DG, the trend toward increased levels of BDNF did not reach statistical significance. Meanwhile a non-significant positive correlation was observed between BDNF gene expression and the number of running wheel counts in APP/PS1 mice treated with exercise, implying that PE can indeed increase

BDNF in these mice. However, APP/PS1 mice treated with BCI-838 and PE displayed indistinguishable levels of BDNF when compared with PE-treated mice, despite the fact that they spent less time on running wheels. This effect shows the potential of BCI-838 to mimic the effects of PE on BDNF expression in the DG. Obviously, modulation of hippocampal levels of BDNF is unlikely to provide a complete explanation for the effects we observed. Unlike P7C3 studied by Choi et al.^29^, BCI-838 stimulated AHN and improved cognition without addition of PE. Thus BCI-838 may provide a pharmacological mimic of PE, at least in its effects on some cognitive function.

Transcriptomic analysis and identification of differential molecular pathways are important steps towards improving our mechanistic understanding and shedding light on the aforementioned unknowns. Our findings further demonstrate that administration of BCI-838 alone induced transcriptomic changes that resulted in up-regulation of BDNF and PIK3C2A of mTOR signaling, down-regulation of IL11RA1 interleukin receptor of AKT signaling, and enriched SP1 and further relevant pathways, implicating neurogenesis and effects of PE. Mechanistically, BCI-838 is considered an orally active ketamine-mimetic, and recent evidence indicates that ketamine modulates mTOR activity and the actions of the physiological effectors of elongation inhibitory factor 4E (eIF4E) and its protein partners, eIF4E binding proteins (eIF4E-BPs). This evidence is also consistent with our findings of dysregulation of a related eukaryotic translation initiation factor, EIF5A. mTOR is known to modulate autophagy^64^. Given the behavioral benefits of BCI-838 in multiple types of proteopathy, its apparent similarity to ketamine regarding mTOR modulation, and the role for mTOR in autophagy, we propose that, like ketamine, BCI-838 exerts its actions through protein translation and autophagy.

Previous studies have explored the effects of exercise or mGluR2/3 modulation alone on AD-related pathology^30, 65^. However, none of these studies explored the effect of the combination of mGluR2/3 modulation and PE in the early stages of AD development in a mouse model. Here we show that BCI-838 improved behavior in APP/PS1 mice and stimulated AHN. Combined with our previous studies in another AD mouse model^30^ as well as a rat model of blast-related traumatic brain injury^31^, three preclinical studies now support the effectiveness of BCI-838 for relief of a range of neurocognitive, neurobehavioral, and neurodegenerative traits. The ability of BCI-838 to recapitulate many of the effects of PE suggests that this and/or other mGluR2/3 antagonists may represent safe and novel pharmacological mimics of PE. If this notion regarding pharmacological mimicry of PE proves to be true, then it will be worth evaluating whether these drugs can prevent, delay, and/or treat neurodegenerative and/or post-traumatic disorders of cognition, anxiety, and/or behavior.

Lifestyle modification remains the only demonstrated disease-modifying intervention for AD (the FINGER study^66^), of which exercise forms a major component, and understanding the action of compounds like BCI-838, which comprise an apparent exercise-related and exercise-independent effect, offers new opportunities to better illuminate these mechanisms, perhaps extend them and inform the development of new therapeutic strategies for this devastating disease.

## Methods

### Animals

The experimental procedures were conducted in accordance with and were approved by the Institutional Animal Care and Use Committee (IACUC) of the James J. Peters VA Medical Center. Studies were conducted in compliance with the US Public Health Service policy on the humane care and use of laboratory animals, the NIH Guide for the Care and Use of Laboratory Animals, and all applicable Federal regulations governing the protection of animals in research. *APP*^*KM670/671NL*^*/ PSEN1*^*Δexon9*^ (APP/PS1) transgenic mice and wild-type (WT) mice were obtained from Jackson Laboratories. Mice were on a C57BL6/J background. All studies used male mice. Three cohorts of mice were used, and the number of animals per group was 20 per iteration.

### Physical exercise (PE) exposure

Cages were fitted with running wheels obtained from Columbus Instruments and equipped with Activity Wheel Monitoring Software. Cages that housed mice in the non-exercise groups were fitted with dummy wheels that the mice could enter but did not rotate. Mice were allowed access to running wheels ad libitum. Running wheel (RW) data (number of wheel revolutions) were recorded.

### Drug administration

BCI-838 was prepared as previously described^30^. Briefly, BCI-838 was dissolved in a solution of 5% carboxymethylcellulose (CMC; Sigma Aldrich) and 0.3% 2N hydrochloric acid solution (Sigma Aldrich) at room temperature. The drug emulsion was prepared daily by sonication for 2 min to allow full dissolution. Animals were divided into five experimental groups: (1) WT mice treated with vehicle (5% CMC); (2) APP/PS1 mice treated with vehicle; (3) APP/PS1 mice treated with vehicle and exposed to running wheels; (4) APP/PS1 mice treated with 5 mg/kg BCI-838; and (5) APP/PS1 mice treated with 5 mg/kg BCI-838 and exposed to running wheels (see Fig. 1). BCI-838 administration was conducted daily between 9 A.M. and 2 P.M. for 28 days by personnel experienced in an oral gavage procedure using methods adapted from Perez Garcia *et al*.^31^. Briefly, mice were firmly grasped and immobilized with the head and body held vertically. A 5-cm straight stainless-steel gavage needle with a 2-mm ball tip (Fischer Scientific) was used for gavage. The gavage needle was wiped clean between animals.

### Bromodeoxyuridine (BrdU) injections

All animals received a single intraperitoneal injection (i.p.) of BrdU (50 mg/kg of body weight) 24 h before sacrifice at the end of the drug treatment. BrdU (Sigma) was dissolved in saline solution (0.9% NaCl in sterile H_2_O) warmed to 40°C and gently vortexed. The solution was allowed to cool to room temperature (25°C) before injection.

### Behavioral testing

Behavioral tests began at the end of the 30 days of drug administration.

### Novel object (NO) recognition

Mice were habituated to the circle arena (30 cm length × 30 cm width × 40 cm height) for 10 min, 24 hours before training. On the training day, two identical objects were placed on opposite ends of the empty arena, and the mouse was allowed to freely explore the objects for 7 min. After 1 h, during which the mouse was held in its home cage, one of the two familiar objects (FOs) was replaced with a NO, and the mouse was allowed to freely explore the familiar and NO for 5 min to assess short-term memory (STM). After 24 h, during which the mouse was held in its home cage, one of the two FOs was replaced with a NO different from the one used during the STM test. The mouse was allowed to freely explore the familiar and NO for 5 min to assess long-term memory (LTM). Raw exploration times for each object were expressed in seconds. Object exploration was defined as sniffing or touching the object with the vibrissae or when the animal!s head was oriented toward the object with the nose placed at <2 cm from the object. All sessions were recorded by video camera (Sentech) and analyzed with ANYMAZE software (San Diego Instruments). In addition, offline analysis by an investigator blind to the treatment status of the animals was performed. Objects to be discriminated were of different size, shape and color, and were made of Lego plastic material. All objects were wiped with 70% ethanol between trials.

### Barnes maze test

The Barnes maze test was performed using a standard apparatus. The testing was conducted in two phases: training (day 1 to 4) and testing (day 5). Before starting each experiment, mice were acclimated to the testing room for 1 h. Mice were transported from their cage to the center of the platform with a closed starting chamber where they remained for 10 sec before exploring the maze. Mice failing to enter the escape box within 4 min on trials 1-4 were guided to the escape box by the experimenter and the latency was recorded as 240 s. Trial 5 was treated as a test trial and mice were given up to 180 s to enter the escape box. The platform and the escape box were wiped with 70% ethanol after each trial. Trials were recorded by video camera and analyzed with ANYMAZE software.

### Tissue processing and immunohistochemistry

Animals were sacrificed at the conclusion of the drug treatment, exercise exposure, and behavioral testing. We induced deep anesthesia with a solution of 100 mg/kg ketamine and 20 mg/kg xylazine, and mice were euthanized by transcardial perfusion with cold PBS (20 ml) and then perfused with 4% paraformaldehyde in PBS. After perfusion, brains were removed and postfixed in 4% paraformaldehyde for 48 h, transferred to PBS, and stored at 4°C until sectioning. Fifty micrometer-thick coronal sections were cut through the entire extent of the hippocampus using a Leica VT1000 S Vibratome (Leica). The sections were stored at -20°C in a cryoprotectant solution (25% ethylene glycol and 25% glycerine in 0.05 M PBS) until processing for immunofluorescence.

Stereology-based counting was performed as described^30^. Every sixth section in a series through the hippocampus was processed for immunohistochemistry so that the interval between sections within a given series was 300 µm. For BrdU staining, DNA was denatured by incubating the sections for 1.5 h in 50% formamide/2× sodium saline citrate (SSC, 0.3 M NaCl and 0.03 M sodium citrate) at 65 °C. Sections were rinsed twice for 5 min in 2× SSC and incubated for 30 min in 2 N HCl at 37 °C. Acid was neutralized in 0.1 M boric acid (pH 8.5). After four 5-min washes with PBS, sections were incubated in blocking buffer (3% goat serum, 0.3% Triton X-100 in PBS) for 1 h and incubated overnight at 4°C with rat anti-BrdU (1:300, Abcam) antibody. The next day, sections were washed four times for 5 min with PBS and incubated for 2 h in the dark with Alexa Fluor 568-conjugated donkey anti-rat IgG (Life Technologies) at a dilution of 1:300. To ascertain the effects of BCI-838, exercise or the combination of treatments on cell proliferation and survival, a second series of sections from each animal was immunolabeled with doublecortin (DCX) and BrdU as described above, using a goat monoclonal anti-DCX antibody (1:500 from Santa Cruz). All slices were mounted onto slides and covered under Fluoro-Gel (with Tris Buffer from Electron Microscopy Sciences).

### RNA extraction and BDNF levels

DG samples were homogenized in QIAzol Lysis Reagent (Qiagen) and total RNA purification was performed with the miRNeasy Micro kit (Qiagen). BDNF mRNA levels were determined by real-time quantitative PCR (qPCR). 200 ng of total RNAs were reversed transcribed using the High-Capacity RNA-to-cDNA Kit (Applied Biosystem). cDNAs were subjected to real-time qPCR in a StepOne Plus system (Applied Biosystem). qPCR in a StepOne Plus system (Applied Biosystem) using the TaqMan Gene Expression Master Mix (Applied Biosystem). qPCR consisted of 40 cycles, 10 s at 95 °C, 20 s at 60 °C, and 15 s at 70 °C each. Ct values were normalized to the expression level of GAPDH. BDNF protein levels were determined similar to the Choi protocol^26^. Briefly, DG tissues from mice of each group were homogenized in RIPA buffer (Sigma, St. Louis, MO) with a cocktail of phosphate inhibitors and then centrifuged at 10,000 x g for 20 min at 4° C. Supernatants were used to measure BDNF using BDNF ELISA kits (R&D Systems, Minneapolis, MN) according to the manufacturer!s instructions. Values were normalized to mg of total proteins.

### Statistical analyses

Values are expressed as mean ± SEM. For behavioral analysis, stereology and spontaneous activity analysis, the statistical tests were performed using the program GraphPad Prism 8.0 (GraphPad Software) or SPSS v26 (IBM). Depending on the behavioral test, multiple comparisons were performed using one-way ANOVA for normally distributed datasets followed by Sidak post hoc tests for multiple comparisons when appropriate or *t*-tests. When repeated-measures ANOVA was used, sphericity was assessed using Mauchly!s test. If the assumption of sphericity was violated (p < 0.05, Mauchly!s test), significance was determined using the Greenhouse-Geisser correction. Pearson correlation coefficients were calculated.

### RNA sequencing

Total RNA from DG of all mice were subjected to RNA sequencing. The sequencing library was prepared according to NEB library prep kit by Novogene.

### Read alignment and gene expression counts

Paired-end RNA-Seq fastq files for 24 samples were aligned to Mouse Reference genome (mm10) using STAR^65^ read aligner. As part of quality control and to allow discrimination between the human APP and PSEN1 transgenes and the mouse App and PSEN1 genes, all samples were also aligned to Homo sapiens Reference genome (Grch38) and corresponding gene expressions were checked for sanity. Mapped reads were summarized to gene counts using subread function of featureCounts^68^.

### Quality control

When variance partitioning was performed to prioritize drivers of variance among our RNA-seq samples using *variancePartition*^69^ R package, both the variation plot of top 50 genes and the violin plot of entire dataset showed a high fraction of unexplained variance (“Residuals”, see Supplemental Data 1). We addressed this by applying surrogate variable analysis (SVA) using *sva*^70^ R package, to estimate unwanted sources of variation. *svaseq* function estimated five surrogate variables (n.sv=5), which were then included in our differential expression analysis as covariates to be accounted for.

### Differential expression analysis

Gene count matrices were generated separately for five groups of 24 samples in total: WT (n=5), APP/PS1 control (n=5), APP/PS1 + BCI-838 (n=5), APP/PS1 + PE (n=4), APP/PS1 combination (n=5); and six primary comparisons: APP/PS1 + BCI-838 vs. APP/PS1 control, APP/PS1 + PE vs. APP/PS1 control, APP/PS1 combination vs. APP/PS1 control, APP/PS1 combination vs. APP/PS1 + BCI-838, APP/PS1 combination vs. APP/PS1 + PE, APP/PS1 + BCI-838 vs. APP/PS1 + PE, and APP/PS1 control vs. WT. The overall gene count matrix was corrected for library size, normalized and resulting 18,357 genes were statistically analysed for DEGs using DESeq2^71^. P-values were adjusted using Benjamini-Hochberg method, where DEGs were defined at a false discovery rate (FDR) of 0.05. Full list of DEGs sorted by their adjusted p-values is provided in Supplemental Data 2.

### Gene set enrichment analysis

DEG sets per primary comparison were tested for statistical enrichment using EnrichR^72^ R package for transcription factors, pathways and gene ontology against 11 relevant public databases. Relevant matches with an FDR of 0.05 were identified. Full list of enrichments is provided in Supplemental Data 3.

### Computational drug repurposing and chemogenomic enrichment analysis

Drug-induced gene expression fold changes were obtained from the Connectivity Map database^43^, which consists of 6,100 individual experiments, representing 1309 unique compounds. The 6,100 individual expression profiles were merged into a single representative signature for the 1,309 unique compounds, according to the prototype-ranked list method^74^. Each compound was scored according to the transcriptomic similarity with BCI-838 transcriptomics (DEGs from APP/PS1 + BCI-838 vs. APP/PS1). Compounds were ranked in order of descending connectivity score. For each compound in the drug signature library, referenced drug–target associations^75^, predicted off-targeting^76, 77^ and side effects were collected. For each of these features, we calculated a running sum enrichment score, reflecting whether that feature was over-represented among the compounds with transcriptomic similarity to BCI-838. Two-tailed p-values were based on comparison with 10,000 permuted null scores, generated from randomized drug target sets that contain an equivalent number of compounds to the true set under evaluation, and adjusted using the Benjamini– Hochberg method. Computational screening and chemogenomic enrichment analysis were performed using R. The full list of drug repurposing results is provided in Supplemental Data 4.

## Acknowledgments

This work was supported by the Department of Veterans Affairs, Veterans Health Administration, Rehabilitation Research and Development Service Awards 1I01RX000684 (SG), 1I01RX002333 (SG), 1I01RX000996 (GE), I01RX002660 (GE), 1I01BX004067 (GE), 1I21RX003459-01 (MAGS), 1I21RX002069-01 (MAGS) and 1I21RX002876-01 (MAGS).

## Author contributions

GPG and MB are co-first authors. CB and FG identified the key pro-neurogenic action of BCI-838 and supplied the drug for these studies; MEE, SG, GPG, JTD designed the study; GPG, JVHM, GMP, AOP performed the experiments; GPG, BR, MB analyzed the data. GPG, MB, SG, MEE, JTD, JVHM, BSG, BR, MS and GE wrote the paper. All authors read and approved the final manuscript.

## Figure legends

**Supplemental Data 1**. Variance partitioning results

**Supplemental Data 2**. Differential gene expression results

**Supplemental Data 3**. Gene enrichment analysis results

**Supplemental Data 4**. Drug repurposing results

