## Supplemental Data 1 for "Orally active mGluR2/3 metabotropic antagonist pro-drug mimics the beneficial effects of physical exercise on neurogenesis, behavior, and exercise-related molecular pathways in an Alzheimer’s disease mouse model"

**Supplemental Data 1.** Variance partitioning results. **a** Variation plot of top 50 genes **b** Violin plot of all genes.

**a.**

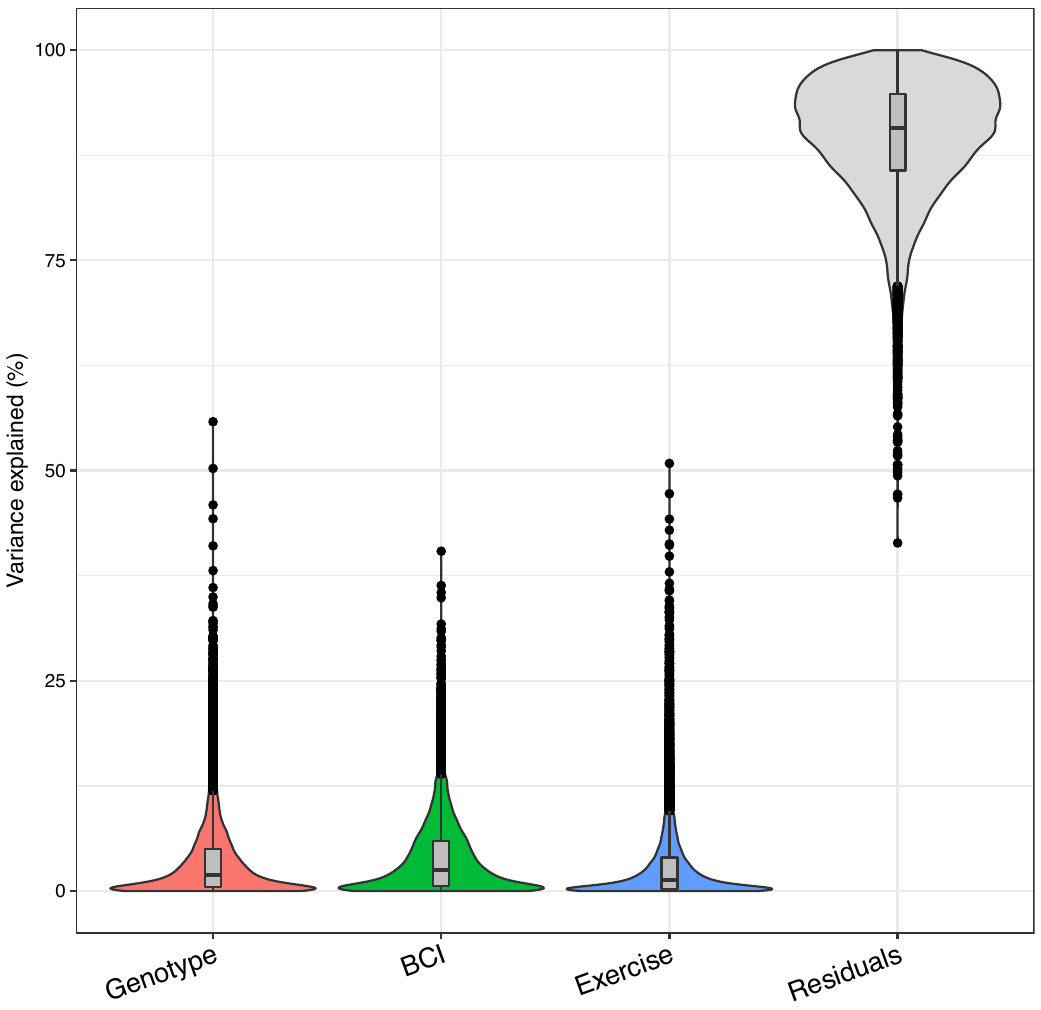

**b.**

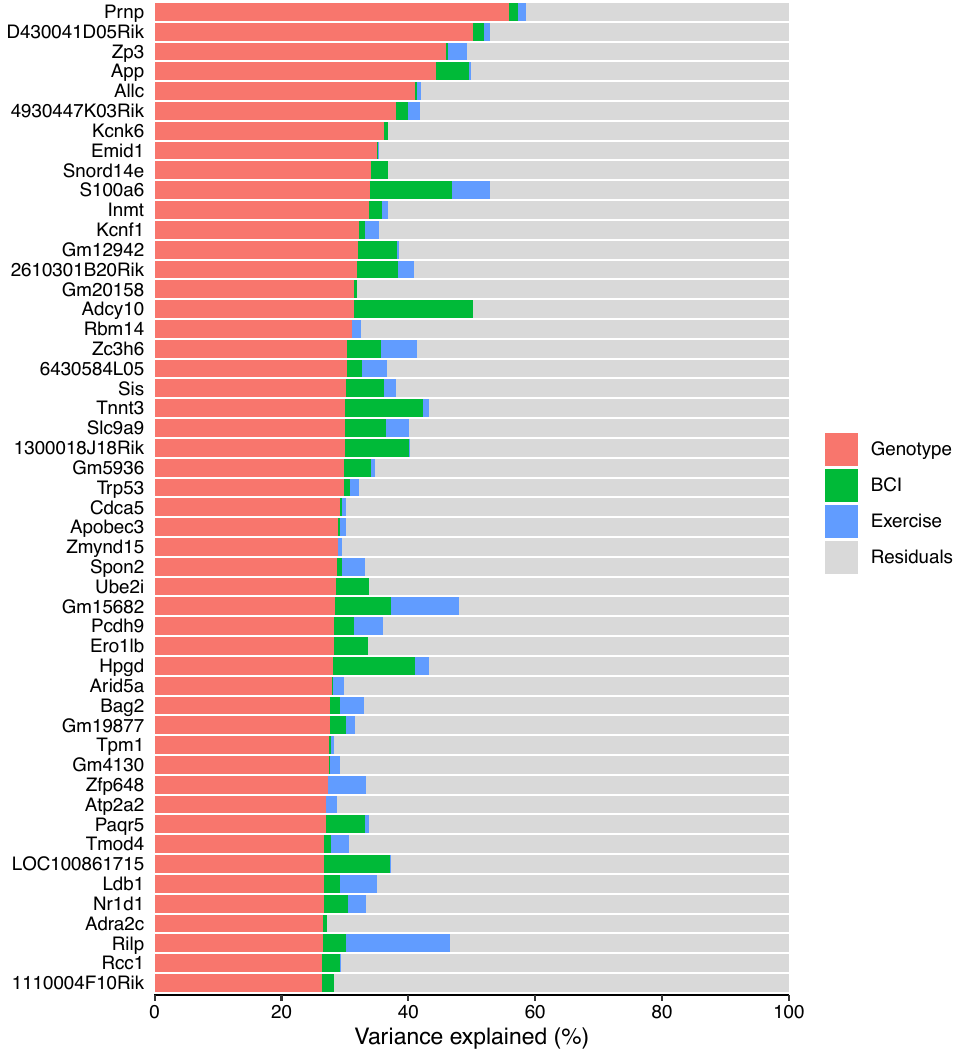
